# Vangl2, a core component of the WNT/PCP pathway, regulates adult hippocampal neurogenesis and age-related decline in cognitive flexibility

**DOI:** 10.1101/2021.01.05.425435

**Authors:** M Koehl, E Ladevèze, M Montcouquiol, DN Abrous

## Abstract

Decline in episodic memory is one of the hallmarks of aging and represents one of the most important health problems facing western societies. A key structure in episodic memory is the hippocampal formation and the dentate gyrus in particular, as the continuous production of new dentate granule neurons in this brain region was found to play a crucial role in memory and in age-related decline in memory. As such, understanding the molecular processes that regulate the relationship between adult neurogenesis and aging of memory function holds great therapeutic potential. Recently, we found that Vang-gogh like 2 (Vangl2), a core component of the planar cell polarity signaling pathway, is enriched in the dentate gyrus of adult mice. In this context, we sought to evaluate the involvement of this effector of the Wnt/PCP pathway in both adult neurogenesis and memory abilities in adult and middle-aged mice. Using a heterozygous mouse model carrying a dominant negative mutation in Vangl2 gene, we show that alteration in Vangl2 expression decreases the survival of adult-born granule cells and advances the onset of decrease in cognitive flexibility. Inability of mutant mice to erase old irrelevant information to the benefit of new relevant ones highlights a key role of Vangl2 in interference-based forgetting. Taken together, our findings show for the first that Vangl2 activity may constitute an interesting target to prevent age-related decline in hippocampal plasticity and memory.

## Introduction

Cognitive decline associated with aging represents one of the most important health problems facing western societies. The memory for specific events, the so-called episodic memory (Eichenbaum, 2000), is particularly sensitive to the effect of aging and declines from early middle-age on (Nyberg and Pudas, 2019). The different processes involved in episodic memory formation, i.e. the ability to encode and retrieve past experiences and to use the learned information in a novel condition, rely upon the hippocampal formation (Eichenbaum et al., 1990; Bunsey and Eichenbaum, 1996), and an impairment of its integrity is considered a key phenomenon in the appearance of age-related memory deficits (Anon, 2016; Gonzalez-Escamilla et al., 2018; Mota et al., 2019).

One key hippocampal sub-structure involved in episodic memory is the dentate gyrus (DG), one of the two brain regions in the adult mammalian brain that retains the capability to produce new neurons throughout adult life. Briefly, the radial glia-like neural stem cells (NSCs) located in the subgranular zone (SGZ), at the interface between the hilus and the granule cell layer (GCL), leave quiescence to proliferate, and through asymmetrical division generate transient amplifying neural progenitor cells (NPCs). These NPCs have the potential to differentiate into neuroblasts and dentate granule neurons (DGN) that mature over several weeks and integrate the existing hippocampal circuitry in order to maintain critical hippocampal functions throughout adulthood (Abrous et al., 2005; Christian et al., 2020). More specifically, rodent studies have shown that adult hippocampal neurogenesis is primarily involved in encoding and retrieving similar events (a process called pattern separation), as well as in using a previously learned information in a flexible way, an ability that is required to navigate through space (Dupret et al., 2008; Baptista and Andrade, 2018). In the context of aging, we have previously uncovered the existence of a link between the rate of new neurons production in the aging DG and memory abilities of senescent animals: preserved memory functions are associated with the maintenance of a relatively high neurogenesis level whereas memory deficits are linked to exhaustion of neurogenesis after learning (Drapeau et al., 2003). More recently, we have deepened this link by showing that the maintenance of healthy newborn neurons in the course of aging provides resilience to cognitive aging (Montaron et al., 2020).

In this context, a better understanding of the molecular processes that regulate the relationship between adult neurogenesis and cognitive aging is important, not only to provide insight on the mechanisms involved in the resilience/vulnerability to cognitive aging but also to develop new preventing/curing strategies. To do so, we focused on the Wnt signaling pathway, because it is compromised in the aging brain (Palomer et al., 2019) and its members were found to regulate different aspects of adult hippocampal neurogenesis (e.g. (Lie et al., 2005; Wexler et al., 2009; Varela-Nallar and Inestrosa, 2013; Mardones et al., 2016; Arredondo et al., 2020)). Recently, we showed that the 4-transmembrane protein Van-Gogh-like 2 (Vangl2) a member of the Wnt core PCP pathway (Montcouquiol et al., 2006b, 2006a), is enriched in young postmitotic neurons of the neurogenic zone of the adult DG (Robert et al., 2020), which led us to investigate the *in vivo* effects of mutation in the Vangl2 gene on adult hippocampal neurogenesis and spatial navigation in the course of aging. We used for this study a mouse model carrying a spontaneous dominant-negative mutation for *Vangl2* (called Loop-tail or Vangl2^Lp^). We show here that mutation of the Vangl2 gene decreases neurogenesis by reducing newborn cells survival. These deficits are associated in adult mice with a dampening in memory flexibility and additional deficits in handling memory interferences and spatial navigation at middle-age. Altogether, this study is the first to show that alterations in the Wnt/PCP signaling pathway is a key factor in accelerating age-related alterations in spatial components of episodic-like memory in association with an impairment of adult neurogenesis.

## Materials and Methods

### Animals

Looptail mutant mice of the LPT/Le stock were originally obtained from Jackson Laboratories (Laboratory stock No. 000220). These mice carry a dominant-negative point mutation at position 464 that causes a serine to asparagine amino acid change resulting in loss of function (Kibar et al., 2001; Murdoch, 2001; Yin et al., 2012). The mouse colony was maintained at Neurocentre Magendie under standard conditions by heterozygous intercross. Only heterozygous *Vangl2^Lp/+^* and their WT littermates *Vangl2^+/+^* were used in this study as homozygous fetuses die perinatally. Heterozygous mice were initially identified by the presence of a looped tail, and biopsies of fixed brains were later analyzed to confirm genotype. Genotyping was performed by direct sequencing of PCR-amplified products generated with the following primers: reverse CTGCAGCCGCATGACGAACT, and forward CCTTCCTGGAGCGATATTTG, designed to flank the point mutation. PCR was performed in 20μL reaction volumes, using GoTaq G2 Hot Start Green Master Mix (Promega), and 0.2μM of each primer. PCR products were sequenced with the forward primer by Sanger method.

Animals were housed in standard plastic rodent cages and maintained on a 12h light/dark cycle (light off at 8pm) with free access to water and food. Only male littermates, housed in individual cages at least 2 weeks before any intervention, were used in all experiments. Two batches of mice were used in order to study on one hand adult neurogenesis (Figure 1), and on the other hand behavioral capabilities (Figure 2, 3 and 4). All procedures and experimental protocols were approved by the Animal Care Committee of Bordeaux (CEEA50) and were conducted in accordance with the European community’s council directive of 24 November 1986 (86/609/ECC).

**Figure 1:**
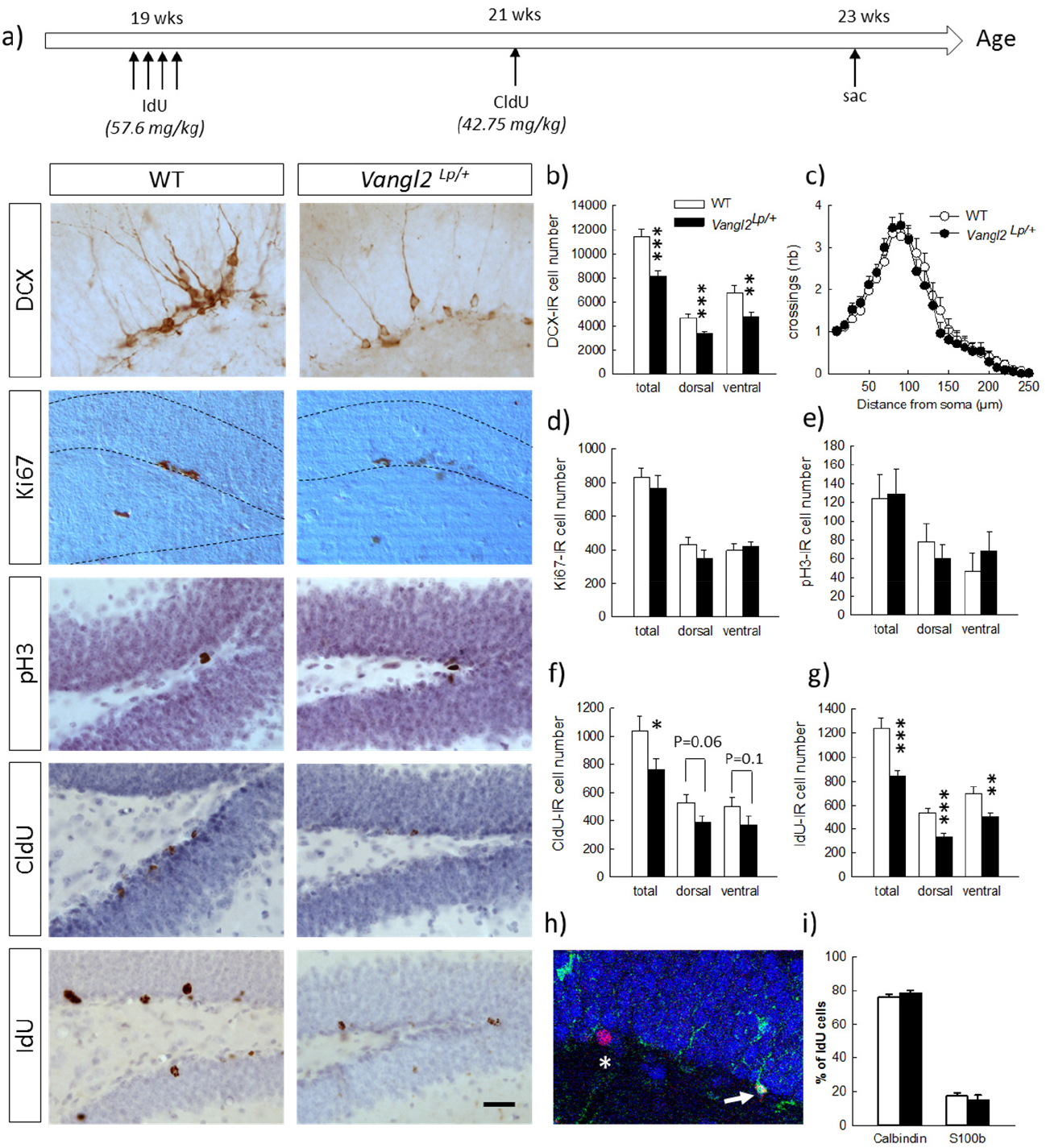
*Vangl2^lp^* mutation decreases survival of adult-born cells in the hippocampal neurogenic niche. a) Timeline of the experiment and representative images of immunostainings in WT and Vangl2^Lp/+^ mice; b) Number of immature neurons (DCX-IR); c) Sholl morphometric analysis of DCX-IR cells; d) Number of Ki67-IR proliferating cells; e) Number of pH3-IR proliferative cells; f) Number of 2-wk old surviving cells (CldU-IR); g) Number of 4-wk old surviving cells (IdU-IR); h) Illustration of a 4-wk old newborn neuron (IdU-Calbindin-IR cell, white star) and a 4-wk old newborn glial cell (IdU-S100-IR cell, white arrow); i) Percentages of neuronal et glial differentiation of 4-wk old IdU-IR cells. White bars, WT n=6-7 mice ; Black bars, Vangl2^Lp/+^ n=8-9 mice; white dots: dendrites WT n=42 neurons; black dots, dendrites Vangl2^Lp/+^ n=32 neurons. * p<0.05; ** p<0.01; *** p<0.001. Scale bar = 50 μm.

**Figure 2:**
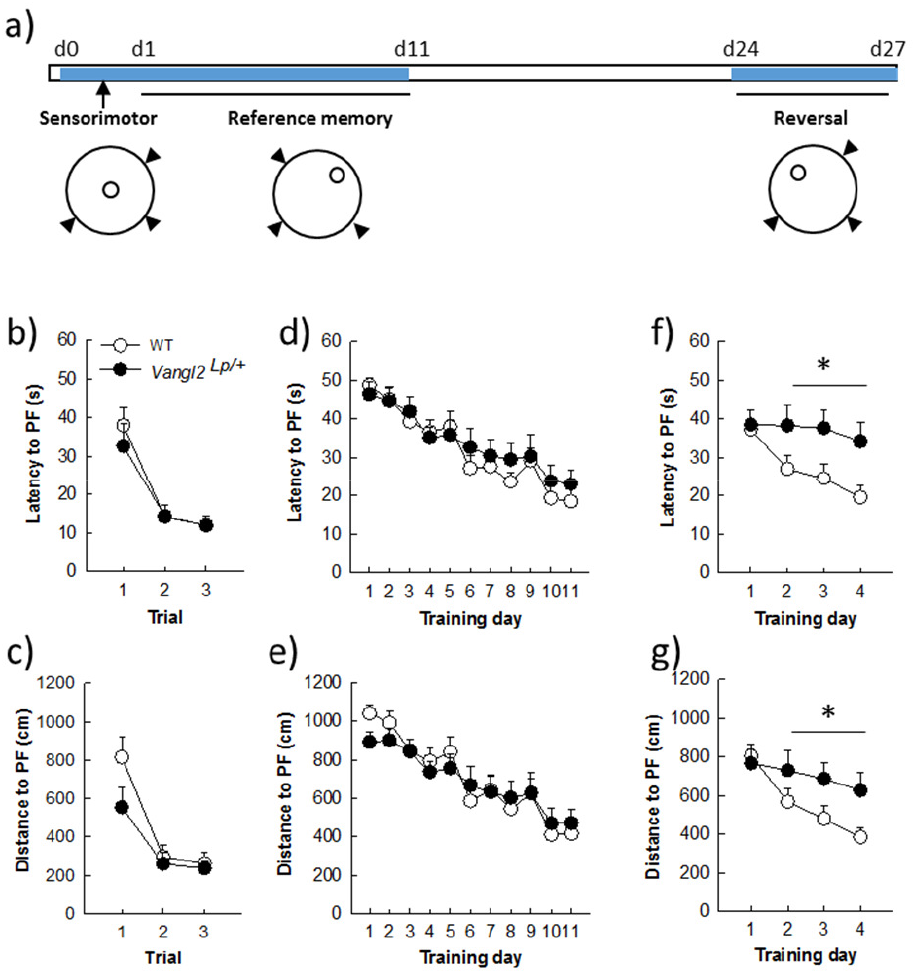
*Vangl2^lp^* mutation leads to deficits in memory flexibility in 4-month old mice. a) Timeline of the experiment. b) Latency to find a cued platform. c) Distance to find a cued platform. d) Latency to find a hidden platform in a reference memory protocol. e) Distance to find a hidden platform in a reference memory protocol. f) Latency to find a hidden platform in a reversal protocol. g) Distance to find a hidden platform in a reversal protocol. White dots, WT n=20 mice ; Black dots, Vangl2^Lp/+^ n=12 mice.

**Figure 3:**
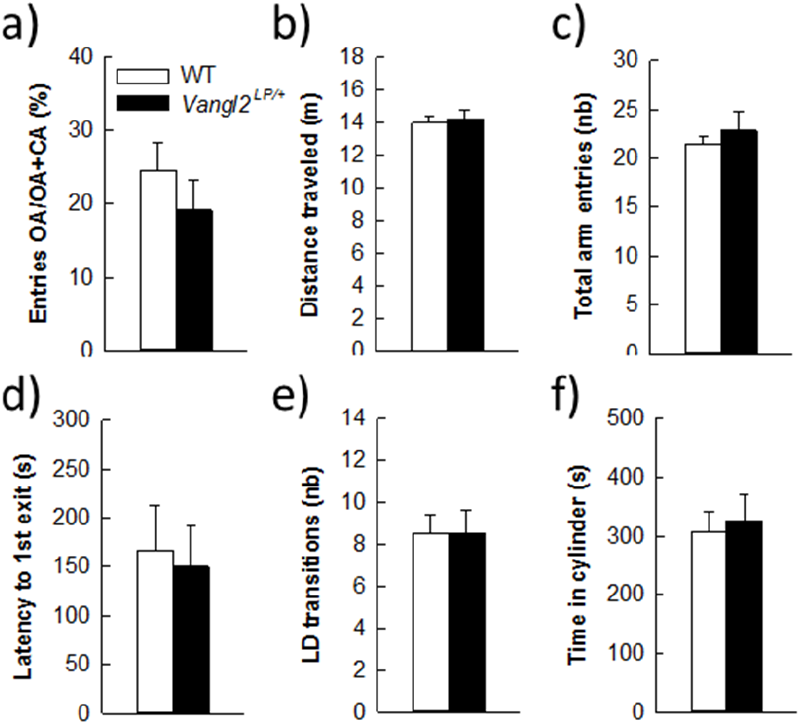
*Vangl2^lp^* mutation does not affect anxiety-related phenotype. Anxiety-like responses were measured in the elevated plus maze (a,b,c) and in the light / dark emergence task (d,e,f). White bars, WT n=21 mice ; Black bars, Vangl2^Lp/+^ n=16 mice.

**Figure 4:**
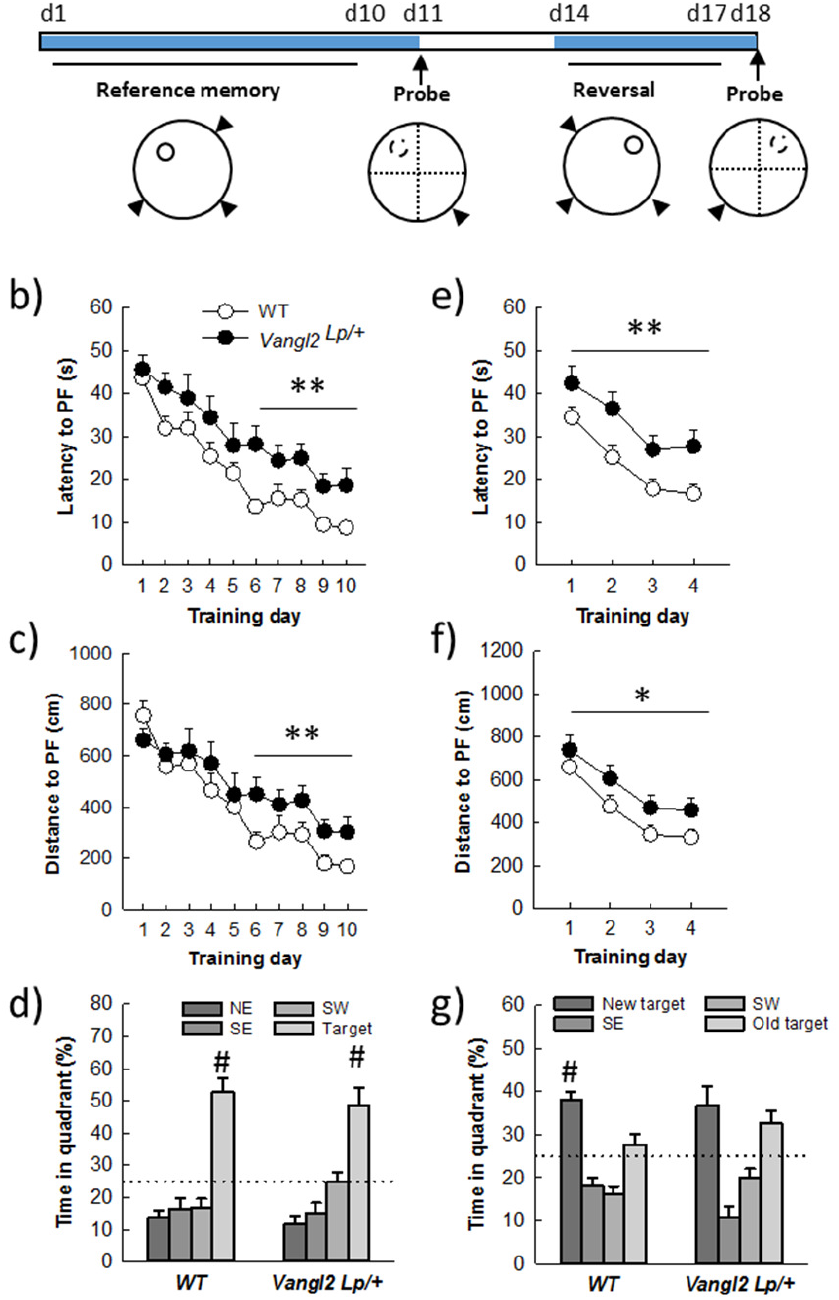
*Vangl2^lp^* mutation increases vulnerability to age-related decline in memory. a) Timeline of the experiment. b) Latency to find a hidden platform in a reference memory protocol. c) Distance to find a hidden platform in a reference memory protocol. d) Percentage of time spent in each quadrant of the watermaze during the probe test. e) Latency to find a hidden platform in a reversal protocol. f) Distance to find a hidden platform in a reversal protocol. g) Percentage of time spent in each quadrant of the watermaze during the second probe test. White dots, WT n=18 mice ; Black dots, Vangl2^Lp/+^ n=11 mice. # different from all other positions, NK p<0.01 for all comparisons.

### Thymidine analogue injections

Naïve 5 month-old mice received a daily i.p. injection of IodoDeoxyUridine (IdU; 57.6 mg/kg dissolved in 1N NH_4_OH/NaCl, Sigma) for 4 consecutive days. Two weeks later, the same mice were injected once i.p. with Chlorodeoxyuridine (CldU; 42.75 mg/kg dissolved in saline, Sigma). Mice were then left undisturbed for 2 weeks upon which they were sacrificed to analyze neurogenesis (Figure 1a).

### Immunohistochemistry

Mice were anesthetized with pentobarbital (100mg/kg) and transcardially perfused with 30 mL of PBS, pH=7.3 and 30 mL of 4% paraformaldehyde in 0.1 M PB, pH=7.3. Brains were extracted and kept in paraformaldehyde until vibratome sectioning. Sections were sequentially collected in 10 sets of serial coronal slices of 40 μm thickness and stored in PBS-azide (0.02%) at 4°C until processing.

Immunoperoxydase detection of each neurogenesis marker was performed on one-in-ten free floating sections. The following primary antibodies were used: mouse anti-BrdU (1/500, BD Biosciences, used for IdU detection), rat anti-BrdU (1/1000, Accurate, used for CldU detection), rabbit anti-Ki67 (1/1000, Novocastra), rabbit anti-pH3 (1/1000, Merck Millipore), goat anti-DCX (1/1000, Santa Cruz biotechnology). Bound antibodies were visualized with horse anti-mouse (1/200, Dako), goat anti-rat (1/1000, Dako), goat anti-rabbit (1/200, Dako), and donkey anti-goat (1/500, Jackson) biotin-labeled secondary antibodies. Briefly, after quenching endogenous tissue peroxidases with 0.3% H_2_O_2_ in methanol for 30 minutes, sections were incubated for 45 min in PBS-triton X-100 (0.3%) in the presence of 3 to 5% of normal serum of goat (for CldU, Ki67, and pH3 stainings), horse (for IdU staining), or donkey (for DCX staining) in order to permeabilize membranes and reduce non-specific binding. Sections were then incubated for 72h at 4°C with primary antibodies. For IdU and CldU stainings, an unmasking step with 2N HCl (30 min at 37°C) was applied prior to primary antibody incubation. Secondary antibody incubation lasted 2h and was followed by avidin-biotin complexes for signal amplification (ABC kit, Dako). Immunoreactivities were visualized using 3,3’-diaminobenzidine as a chromogen. Finally IdU and CldU stained sections were counterstained with hematoxylin in order to analyze DG and OB volumes.

For phenotyping newborn cells, triple immunofluorescent IdU-Calbindin-S100 labeling was performed; floating sections were first treated with 2N HCl (30 min at 37°C), incubated for 45 min in PBS containing 5% goat normal serum and 0.3% triton-X-100, followed by 72h of incubation with a mixture of mouse anti-IdU (1/1000; BD biosciences), guinea pig anti-calbindin (1/300, Synaptic System) and rabbit anti-S100 (1/1000, Sigma) antibodies in PBS-Triton-X-100. Immunoreactivities were revealed with Alexa 568 goat anti-mouse (1/1000, Invitrogen), Alexa 647 goat anti-guinea-pig (1/1000, Jackson), and Alexa 488 goat anti-rabbit (1/1000, Invitrogen) secondary antibodies. Sections were mounted on glass slides and coverslipped with polyvinyl alcohol mounting medium with 1,4-diazabicyclo[2.2.2] octane (PVA-DABCO).

### Stereological analysis

An unbiased estimate of IdU, CldU, Ki67, pH3, and DCX -positive cell number in the dentate gyrus was calculated using every tenth section along the rostro-caudal axis. Exhaustive counting in the left hemisphere under 630x or 1000x magnification was used for all markers. Dorsal sections of the DG were defined as sections with coordinates extending from −1 to −2.4 mm antero-posterior from Bregma, and ventral sections included sections from −2.4 to −3.9 from Bregma. For the olfactory bulb (OB), 50×50 mm counting frames at evenly spaced intervals of 300×300 mm, with an exclusion guard zone of 2 mm, was used to evaluate the number of IdU-IR cells (Stereo Investigator software, MicroBrightField, VT, USA). Results are expressed in the DG as the total number of cells in both hemispheres, and in the OB as cell volumetric density (number of cells per mm^3^).

### Dendrite analysis

Dendritic processes of DCX-IR cells were traced using NeuroLucida (MicroBrightField, Williston, Vermont). Only neurons showing tertiary processes with a vertically-oriented cell body, a complete staining of the dendritic tree and that were not overlapping with neighboring cells were selected for analysis. Soma size, number of branch points, and total length of the immunopositive dendritic tree were measured. A Sholl analysis was further conducted on the reconstructed neurons.

### Phenotype analysis

The percentage of IdU-labeled cells co-expressing calbindin or S100 was determined throughout the DG. For each animal, IdU-positive cells were randomly selected and analyzed for coexpression with calbindin or S100. Using a confocal microscope (DMR TCS SP2; Leica Microsystems) equipped with a 63 PL APO oil objective, an argon laser (488 nm), a green helium-neon laser (543 nm) and a red helium-neon laser (633nm) each selected IdU-positive cell was analyzed in its entire z axis using 1 μm intervals.

### Behavioral testing

A separate batch of mice was tested for behavioral capabilities. Naïve 3 month-old mice were first tested for anxiety-related behavior. One month and 9 months later (at 4 and 12 months of age), their learning and memory abilities were measured in the Morris Water maze.

### Measurement of anxiety-related behaviors

#### The elevated plus maze (EPM)

was conducted in an apparatus composed of transparent Plexiglas with two open (45 x 5 cm) and two enclosed (45 x 5 x 17cm) arms that extended from a common central squared platform (5 x 5 cm). The floor of the maze was covered with black makrolon and was elevated 116 cm above the floor. A small raised lip (0.5 cm) around the edges of the open arms prevented animals from slipping off. The test session began with the mouse individually placed on the center square facing an open arm. Animals were allowed to freely explore the maze for 5 min under mildly anxiogenic conditions (dim light of 45 lux using a halogen lamp). A camera connected to a computer was utilized to track the mouse path during the entire session (©VideoTrack, Viewpoint). Automatic path analysis measured the time spent in and the total number of entries into the open and closed arms.

#### The light/dark emergence test (LD)

was conducted 2 days later in an open-field (square arena of 50 × 50 cm closed by a wall of 50 cm high and made in white PVC) brightly lit (~400 lux) containing a cylinder (10 cm deep, 6.5 cm in diameter, dark gray PVC) located length-wise along one wall, with the open end 10 cm from the corner. Mice were placed into the cylinder and their behavior recorded for 15 min with a videotracking system (©VideoTrack, Viewpoint). Initial latency to emerge from the cylinder, defined as placement of all four paws into the open-field, as well as total number of entries into the cylinder and total time spent inside the cylinder were analyzed.

### Measurement of learning and memory abilities in the water maze

Testing took place in a circular swimming pool (150 cm in diameter) located in a room with various distal cues, and filled with water maintained at room temperature (19-20°C) and made opaque by the addition of a nontoxic cosmetic adjuvant. Data were collected using a video camera fixed to the ceiling of the room and connected to a computerized tracking system (Videotrack, Viewpoint) located in an adjacent room that also contained the individual home cages of the mice during testing. The tracking system allowed calculation of escape latency and path length covered by a mouse until it finds the platform.

#### Pre-training

mice received a two-step pre-training session. First, they were placed for 15 sec onto an escape platform (14 cm diameter, 1 cm above water surface) located in the center of the pool upon which they were released into the water for 30 sec and guided to the platform where they had to stay for 15 sec. This step was repeated until all mice were able to climb and stay onto the platform for 15 sec (usually 1 trial). Once all mice had acquired this step, the platform was lowered below water surface (1 to 1.5 cm) and the same procedure was applied until all mice were able to stay onto the platform for 15 sec (usually 3 to 4 trials).

#### Sensorimotor testing

The next day sensory motor abilities of all mice were tested during three trials with a 60s cut-off and a 5 min inter-trial interval (ITI) using a visible cued platform located in the northwest (NW) quadrant of the pool. A trial terminated when the animal climbed onto and stayed on the platform for 15 sec. Mice that failed to find the platform within a 60 second cut-off time were placed on it for 15 sec by the experimenter. Between each trial, mice were held in their home cages. For both pretraining and sensorimotor testing, the pool was surrounded by a black curtain with no cues. For the rest of the experiment, the curtain was removed and mice had access to the distal cues located in the room.

#### Training with variable start positions

On the day following sensorimotor testing, mice were trained for 3 daily trials with a cut-off of 60 sec and 5 min ITI to find the platform located in the northeast (NE) quadrant. Mice were released from 3 different starting points at each trial, and different sequences of starting points were used day by day. Training lasted 11 days.

#### Reversal

Thirteen days after the last session, mice were trained in a reversal protocol whereby they had to find the new location of the platform that was moved to the northwest (NW) quadrant. Their performances were analyzed over 4 days using 3 daily trials with a 5 min ITI.

#### Training in middle-aged mice

Eight months later, mice were tested again in the same task using the same procedures (two WT and 1 heterozygous mice had to be eliminated for inconsistent behavior): they were first exposed to 10 days of training to find the hidden platform located in the NW quadrant, upon which a probe test was performed to measure memory of the platform location; to this end, the hidden platform was removed from the pool and each subject was placed into the water diagonally opposite to the target quadrant. Time spent in the target quadrant (% of total time, chance level = 25%) was measured over 60 sec.

Two days later their ability to locate the same platform moved to the NE quadrant was also assessed over 4 days in a reversal test. Twenty four hours later, a probe test was performed.

### Statistical analyses

All statistical analyses were performed with Statistica 12.0 software (Statsoft). Neurogenesis and anxiety-related data were analyzed with Student’s *t* tests, while water maze data were analyzed with two-way ANOVAs using NK post hoc test whenever appropriate. All data are presented as mean +/− SEM.

## Results

### Vangl2^Lp^ mutation affects the survival of adult-born neurons

First, we investigated the impact of the Looptail mutation on the number of cells immunoreactive for doublecortin (figure 1b), an immature neuronal marker used as a surrogate of neurogenesis. We found a decreased number of DCX-IR cells in both the dorsal (t_14_=-3.53, p=0.003) and ventral dentate gyrus (DG) (t_14_=-2.67, p=0.01) resulting in an overall decrease (t_14_=-4.14, p=0.0009) and indicating that looptail mutation decreases the number of cells engaged in a neuronal lineage. We analyzed the dendritic morphology of these newborn cells and performed Sholl analyses on DCX-IR cells with tertiary dendrites. We found that neither cell body area (t_72_=-0.56, p=0.57), number of branching points (t_72_=-0.62, p=0.53), dendritic length (t_72_=0.58, p=0.56), or distribution of dendrites along the dendritic tree (figure 1c; genotype effect F_1,71_=0.09, p=0.76; genotype x distance from soma interaction F_1,24_=0.67 p=0.87) were influenced by the Vangl2^Lp^ mutation. In order to determine whether the reduction in DCX-IR cells was linked to a deficit in cell proliferation and/or a deficit in cell survival, we first labeled dividing cells using Ki67 and phosphorylated-Histone 3 (pH3). We found no differences in the number of cells immunoreactive for Ki67 or pH3 between WT and *Vangl2^Lp/+^* mice whether in the dorsal (t_13_=-1.18 p=0.25; t_12_=-0.71 p=0.48, for Ki67 and pH3, respectively), ventral (t_13_=0.53 p=0.60; t_12_=0.79 p=0.44, for Ki67 and pH3, respectively), or total DG (t_13_=-0.66 p=0.52; t_12_=-0.12 p=0.90, for Ki67 and pH3, respectively), indicating that looptail mutation does not affect cell proliferation (figure 1d, 1e). To analyze cell survival, mice were injected with the thymidine analogs IdU and CldU at 2 weeks interval and the number of surviving cells was counted 2 weeks after the last injection (figure 1a). Two-week old CldU-IR cell number (figure 1f) was reduced in *Vangl2^Lp/+^* mice compared to WT (t_14_=-2.06, p=0.05), with a contribution of both the dorsal and ventral parts, although differences did not reach significance (t_14_=-1.99 p=0.06 and t_14_=-1.49 p=0.15 for dorsal and ventral DG, respectively). This reduced survival of newly-born cells was confirmed in 4 weeks old IdU-IR cells (figure 1f; t_13_=-3.97, p=0.001) in both the dorsal (t_13_=-4.11, p=0.001) and ventral DG (t_13_=-2.84, p=0.01). This was not accompanied by any modification in the volume of the granule cell layer (t_14_=-0.72, p=0.47). If altogether these data indicate an involvement of Vangl2 in cell survival, the larger effect observed in IdU-IR cells suggests a mechanism targeting preferentially 2-4 weeks old cells. We thus analyzed whether the fate of these surviving cells was affected by the looptail mutation (figure 1h, 1i) and found no differences in the percentage of 4-wk old cells acquiring a neuronal (IdU-Calbindin-IR cells, t_12_=0.80, p=0.43) or a glial phenotype (IdU-S100-IR cells, t_12_=-0.64, p=0.52). Together with the reduced number of IdU-IR cells, these data indicate a decreased production of newborn neurons in *Vangl2^Lp/+^* mice, and support a role for Vangl2 in adult-born cell survival.

To determine whether this role is specific for the DG, we measured cell survival in the OB and found no difference in the volume of the GCL (t_13_=-0.72, p=0.47), or the number of 4-wk old IdU—IR cells (WT n=65545 +/− 5104, *Vangl2^Lp/+^* n=61651 +/− 1916; t_13_=-0.75, p=0.46) between WT and *Vangl2^Lp/+^* mice. Although these results do not preclude a specific effect on other parameters of adult neurogenesis, such as neuroblasts migration or neuronal morphology, which were found to be altered in the OB after knocking-down Vangl2 expression using constructs with dominant-negative forms (Hirota et al., 2012), they nevertheless indicate that Vangl2 involvement in neuronal survival is specific to the hippocampal neurogenic niche.

### Vangl2^lp^ mutation alters memory flexibility but not anxiety-related behavior in adult mice

We next sought to investigate the consequences of these neurogenesis deficits on mice abilities to navigate through space. Because of their phenotype (looptail), we first tested whether mutant mice were impaired in their abilities to swim toward a goal and exposed them to 3 trials with a cued platform (figure 2a). Although slightly less rapid than WT mice (mean swim speed WT=21.81 +/− 0.34 cm.s^-1^, *Vangl2^Lp/+^* = 18.03 +/− 0.7 cm.s^-1^; genotype effect F_1,30_=27.98, p<0.0001) heterozygous mice did not significantly differ in their latency (figure 2b; genotype effect F_1,30_=0.23, p=0.63; genotype x trial interaction F_2,60_=0.34, p=0.71) or distance (figure 2c; genotype effect F_1,30_=2.61, p=0.11; genotype x trial interaction F_2,60_=1.42, p=0.25) to reach the platform over the 3 trials. Then animals were tested for their ability to find a hidden platform (the position of which remained unchanged during training) from variable starting positions. This type of memory, i.e. reference memory was not affected either and both latency (Figure 2d) and distance (figure 2e) to find the platform decreased similarly in *Vangl2^Lp/+^* and WT mice over days of training (latency: genotype effect F_1,30_=0.35, p=0.55; genotype x training day interaction F_10,300_=0.48, p=0.89; distance: genotype effect F_1,30_=0.04, p=0.83; genotype x training day interaction F_10,300_= 0.68, p=0.74).

Given that we and others have shown that new neurons are involved in cognitive flexibility (Dupret et al., 2008; Garthe et al., 2009), mice were then tested in a reversal version of the test: the position of the platform was changed and the ability of mice to learn the new position (and forget the old one) was measured. This procedure revealed major deficits in *Vangl2^Lp/+^* mice, with an increase in both latency (figure 2f) and distance (figure 2g) to find the new platform location, although the difference did not reach significance for distance (latency: genotype effect F_1,30_=5.16, p=0.03; genotype x training day interaction F_3,90_=2.06, p=0.11; distance: genotype effect F_1,30_=3.14, p=0.08; genotype x training day interaction F_3,90_=2.41, p=0.07). When analysis was performed over the last 3 days of training, which reflect the actual learning phase of the new platform location, both latency and distance were significantly altered in *Vangl2^Lp/+^* mice (latency: genotype effect F_1,30_=7.06, p=0.01; genotype x training day interaction F_2,60_=0.16, p=0.84; distance: genotype effect F_1,30_=5.37, p=0.02; genotype x training day interaction F_2,60_=0.25, p=0.78) reflecting a failure to adapt to the changed situation and to develop appropriate new spatial preference.

To expand these data and control the emotional status of mice, in light of neurogenesis’ involvement in anxiety (Revest et al., 2009), we tested whether *Vangl2^Lp/+^* mice displayed any behavioral irregularities in two tests aimed at measuring their anxiety level. In the elevated plus maze where anxious mice usually avoid open arms, WT and heterozygous mice spent a similar amount of time in these arms (figure 3a; t_35_=-0.97, p=0.33) and showed the same level of general exploration (total distance travelled: figure 3b, t_35_=0.27, p=0.78; total number of visited arms: figure 3c, t_35_=0.70, p=0.48). When given the choice between an aversive brightly lit and open environment and a typically preferred dark and enclosed one in the Light/Dark test, mice from both genotypes took the same time to engage into exploring the light environment (figure 3d, t_35_=-0.27, p=0.78), made the same number of transitions between the 2 compartments (figure 3e, t_35_=0.02, p=0.97) and spent a similar amount of time in the enclosed reassuring environment (figure 3f, t_35_=0.34, p=0.73). Together these results demonstrate that looptail mutation is not a sufficient condition to impact the anxiety level of mice.

### Vangl2^Lp^ mutation alters capability to handle memory interferences in middle-aged mice

We next questioned whether the deficit in flexibility observed in adult mice would persist and whether memory abilities worsen with age. At 12 months of age, the same mice were retested in the water maze (figure 4a). Twelve months old WT mice learned more quickly the task compared to 4 months old (age effect on latency: F_1,16_=38.56, p<0.001), while *Vangl2^Lp/+^* mice showed the same rate of learning (age effect on latency: F_1,10_=2.76, p=0.12) indicating that they were not sensitive to the beneficial effect of retraining (genotype x age interaction on latency:F_1,26_=4.56, p=0.04; WT 12-month <WT 4-month at p<0.001; Vangl2 ^Lp/+^ 12-month = 4-month). As a result, analysis of learning curves in 12 months old mice revealed a delay in learning platform location over days in *Vangl2^Lp/+^* mice compared to WT as revealed by differences in latency (figure 4b, genotype effect F_1,27_=9.15, p=0.005; genotype x training day interaction F_9,243_=0.61, p=0.78). Learning was then divided into 2 phases: an early phase (1^st^ 5 days) during which there is a fast and large improvement in performance and a late phase (last 5 days) during which performances remain at a stable level (Dobrossy et al., 2003). We found that *Vangl2^Lp/+^* mice were impaired during the last phase of learning, indicating deficits in memory stabilization (latency: genotype effect F_1,27_=13.36, p=0.001; genotype x training day interaction F_4,108_=0.54, p=0.70; distance: genotype effect F_1,27_=8.01, p=0.008; genotype x training day interaction F_4,108_=0.24, p=0.91). This delay did not affect heterozygous mice memory for platform location tested 24h after the last training trial in a probe test as mice from both genotypes spent more time in the target quadrant (TQ, NorthWest position) than in any other one (WT: quadrant effect F_3,51_=25.82, p<0.0001; TQ > each other quadrant at p<0.001; *Vangl2^Lp/+^:* quadrant effect F_3,30_=14.43, p<0.0001; TQ > each other quadrant at p<0.001). We next carried out a reversal test and found that heterozygous mice were impaired in their rate of learning as both latency and distance to reach the platform were increased compared to WT (latency: figure 4e, genotype effect F_1,27_=11.26 p=0.002; genotype x training day interaction F_3,81_=0.18 p=0.90; distance: figure 4f, genotype effect F_1,27_=4.49 p=0.04; genotype x training day interaction F_3,81_=0.13 p=0.93). A probe test performed after 4 days of training indicated that while WT mice acquired the new platform location and spent significantly more time in the new target quadrant (new TQ) than in any other one (quadrant effect: F_3,51_=19.12, p<0.0001; new TQ > each other quadrant at p<0.01), *Vangl2^Lp/+^* mice persevered swimming to the previous platform position (quadrant effect F_3,30_=10.66, p<0.0001, new TQ = old TQ > SE and SW at p<0.05) confirming deficit in cognitive flexibility. Altogether these data indicate that Vangl2 may be involved in protecting cognitive capabilities from age-related decay.

## Discussion

In this study we investigated the involvement of the core PCP gene *Vangl2* in adult hippocampal neurogenesis and spatial memory abilities in the course of aging. We report that knocking-down Vangl2 expression in looptail mice is a sufficient condition to impair the survival of adult hippocampal newborn neurons and alter cognitive flexibility in adult mice. In addition it accelerates age-related deterioration of episodic-like memory by increasing susceptibility to interference, leading to deficits in spatial navigation. Altogether these results demonstrate for the first time a key role for members of the Wnt/PCP pathway in regulating neurogenesis-dependent episodic-like memory in the course of aging.

### Looptail mutation regulates the number of adult-born neurons

Although the PCP pathway has been extensively studied in embryonic development, the consequences of its defects in adults are poorly understood. Given the known role of this pathway in actin cytoskeleton remodeling and cell migration, we hypothesized that the PCP pathway might be important for several aspects of adult neurogenesis. We indeed report a decrease in the number of newly-born neurons resulting from a diminution of the survival of 2- to 4-wk old newborn cells, without modifications in cell proliferation. This result is in line with the few studies that have analyzed the involvement of PCP components on hippocampal adult neurogenesis. Thus Wnt-5a, which directly regulates the level of Vangl2 phosphorylation through Ror2, a step required for Vangl2 function in PCP during mammalian development (Gao et al., 2011; Yang et al., 2017), reduces the number of DCX-IR cells and the number of newborn neurons reaching maturity without modifying cell proliferation (Arredondo et al., 2020). The lack of effect on cell fate determination is also consistent with the consequences of knocking down FZD3 or Celsr1-3, other core PCP components (Schafer et al., 2015). Although the overall data available are thus still too sparse to draw mechanistic hypothesis, our observations may result from both a direct effect of Vangl2 altered expression in DCX-IR cells and an indirect effect of disturbances in the cellular niche that encompasses developing newborn neurons. Interestingly, whatever the involved mechanisms, neurogenesis in the OB was spared, indicating a different role of Vangl2 in the two adult neurogenic niches.

These effects are clearly distinct from those reported for the canonical members of the Wnt pathway, which have been proposed to be important for regulating the balance between maintenance of the stem cell pool and differentiation of newborn neurons (Wexler et al., 2009; Varela-Nallar and Inestrosa, 2013) based on the following evidences: i) general blockade of Wnt signaling in the DG reduces proliferation and neurogenesis while its activation increases neurogenesis (Lie et al., 2005); ii) the Wnt signaling inhibitors Dickkopf 1 (Dkk1) and secreted Frizzled-related protein 3 (sFRP3) negatively regulate neurogenesis (Jang et al., 2013; Seib et al., 2013; Sun et al., 2015); iii) Frizzled 1 (FZD1), a well-known receptor for the Wnt/β-catenin pathway, controls neuronal differentiation and newborn cells migration (Mardones et al., 2016); iv) manipulation of GSK-3β levels, a key player in Wnt/β-catenin activation, affects proliferation and neural fate specification (Eom and Jope, 2009; Fiorentini et al., 2010).

Together with our data, this suggests a complementary role for canonical and PCP Wnt components in regulating different aspects of adult neurogenesis.

Because one of the main target of core PCP signaling is the modulation of the cytoskeleton (Babayeva et al., 2011), we expected that Vangl2 would regulate the branching of dendritic trees in newborn neurons. This would have been in line with recent studies in which downregulation of Vangl2 using shRNA, alteration in Vangl2 expression in mice carrying a null mutation of Vangl2, or in heterozygous looptail mice, led to a global decrease in the complexity of dendritic branching and the number of spines in dissociated hippocampal and cortical neurons (Hagiwara et al., 2014; Okerlund et al., 2016). In accordance with these results obtained in cultured cells, other *in vivo* studies reported a similar reduction in dendritic tree complexity and spine density specifically in adult-born granule cells after disruption of other PCP genes such as Wnt-5a, FZD3 or Celsr2/3 (but not Celsr1) in the adult DG (Schafer et al., 2015; Arredondo et al., 2020). However, these results are contrasted by a recent study using conditional deletion of Vangl2 in which an increase in spine density of hippocampal pyramidal cells and an opposite role for Vangl2 and Celsr3 were reported (Thakar et al., 2017). Altogether, these data suggest a complex role of Vangl2 in regulating neuronal morphology and spinogenesis that has been further emphasized in our recent *in vivo* study showing that postnatal deletion of Vangl2 leads to different alterations of the dendritic arbor and spine density of Golgi-stained dentate granule neurons (DGNs) depending on distance from soma, with decreased complexity in the proximal section and increased ramification in the more distal one (Robert et al., 2020). Interestingly, we also reported in this study a dual distribution of Vangl2 protein during the maturation period of granule cells, with an enrichment in the cell bodies of immature postmitotic DGNs and massive redistribution to the neurites of mature DGNs (Robert et al., 2020). This may lead to an enrichment of Vangl2 in the cell bodies of DCX-IR cells that could explain why dendritic branching was not affected in our targeted population, although differences in experimental approaches and models (spontaneous mutation versus transgenic mice and inducible conditional KO) may also certainly explain these divergent results.

### Looptail mutation accelerates age-related memory disorders

The consequences of these alterations on spatial navigation were studied in an open space using the water maze in which animals learn the location of a hidden platform using distal cues, the starting point being changed at each trial. This task requires a high degree of flexibility as mice have to use an allocentric mapping strategy that consists in learning the positional relationships among multiple independent environmental cues (“spatial relational memory’’). This relational representation is needed to support the flexible use of learned discriminative cues in novel situations (i.e. changing starting position), and is consequently necessary to solve the task. Given that we could not evidence any behavioral perturbation using this protocol in 4-month-old Vangl2 male mice, task difficulty was increased by changing the goal’s spatial location (platform position). This procedure, called reversal, was sufficient to uncover a deficit in flexibility in mutant adult mice as they were unable to adapt their strategy. This deficit was also evident when mice were retested at middle age, indicating that the early deficit of Vangl2 is consistent with an advanced onset of a decrease in flexibility otherwise observed with normal aging in mice (Matzel et al., 2011; Yang et al., 2019), rats (Mota et al., 2019), non-human primates (Joly et al., 2014) and humans (van Boxtel et al., 1998). It is also interesting to highlight that although repeated training improved performances in control mice, mutant mice did not benefit from this previous training suggesting that they did not remember the task. In fact memory deficits worsened with age as middle-aged mutant mice were impaired during both the initial learning phase and during reversal. When a probe test was performed at the end of the reversal, we observed that *Vangl2^Lp/+^* mice persevered swimming to the previous platform position confirming deficits in cognitive flexibility that may result from their inability to erase old irrelevant information to the benefit of new relevant ones. In their outstanding review, Davis and Zhong (2017) described different mechanisms of active forgetting (Davis and Zhong, 2017), and although we favor an alteration of interference-based forgetting in *Vangl2^Lp/+^* mice, an alteration of retrieval-induced forgetting and/or intrinsic forgetting can also be at play. Finally, although the present dataset suggests that adult neurogenesis mediates the involvement of Vangl2 in forgetting, it does not exclude the participation of others mechanisms involving downstream effectors such as RAC1 (Liu et al., 2016) and/or AMPA signaling (Robert et al., 2020) (for review (Davis and Zhong, 2017)).

Altogether our results are consistent with the putative role of adult-born neurons in relational memory, flexibility and active forgetting. For instance, regarding flexibility, we reported that specifically depleting adult neurogenesis led to an incapacity for mice to find the platform location when they were released from a novel starting point after acquiring the location of this platform using constant starting points (Dupret et al., 2008). Regarding active forgetting, several recent studies strongly support a role for adult newborn neurons (Akers et al., 2014; Epp et al., 2016; Tran et al., 2019). In particular, computational and experimental models taken together indicate that decreasing neurogenesis stabilizes existing memories, thus enhancing interference for the encoding of new memories, particularly when existing and new memories overlap in content, as is the case with goal reversal in the water maze (Aimone et al., 2009; Epp et al., 2016). As a consequence, this impedes the encoding of the new, conflicting information, as we observed in mutant mice. Although such stabilization may hold an adaptive value under physiological conditions, in the present case, it reflects an inability to adjust to the changing environment and thus a lack of flexibility.

Finally, we propose that the decreased neurogenesis observed from early adult life on in mutant mice is involved in the accelerated aging of their cognitive functions. Indeed, we and others have previously shown that adult neurogenesis is one of the core mechanisms in the aging of memory functions (Drapeau and Abrous, 2008; McAvoy and Sahay, 2017). For instance we have shown that increasing and decreasing adult neurogenesis through modulation of the HPA axis activity protects and prevents, respectively, from the appearance of memory disorders measured in the Water maze from middle-age on (Lemaire et al., 2000; Montaron et al., 2006). Using more specific approaches, recent studies have confirmed this link, showing that enhancement of adult-born DGCs integration by genetic overexpression of Kruppel-like factor 9, a negative regulator of dendritic spines, in mature dentate granule neurons (McAvoy et al., 2016) or enhancement of neural stem cells expansion by overexpression of the cell cycle regulators Cdk4/cyclinD1 (Berdugo-Vega et al., 2020) improve memory abilities of middle-aged mice.

In summary, the results presented here show for the first time that a single allele deregulation of Vangl2 is sufficient to impair adult hippocampal neurogenesis and cognitive flexibility when animals are young adults and to accelerate age-related decline in spatial learning most probably through an alteration of interferences-induced forgetting. These results pinpoint that Vangl2 activity could be a predictive factor of successful aging and open tantalizing opportunities for the prevention of age-related cognitive deficits.

## Acknowledgment

We are grateful to Miss D. Gonzales and the staff of the Animal Housing and Genotyping facilities funded by Inserm and LabEX BRAIN ANR-10-LABX-43 for mouse genotyping. We thank Miss F. Martell and Mr C. Dupuy for providing excellent animal care. This work was supported by Institut National de la Santé et de la Recherche Médicale (to DNA), ANR MemoNeuro ANR2010-BLAN-1408-01 (to DNA) and the Aquitaine Region (to DNA).

## REFERENCES

Abrous DN, Koehl M, Le Moal M (2005) Adult neurogenesis: from precursors to network and physiology. Physiol Rev 85:523–569.

Ageta H, Murayama A, Migishima R, Kida S, Tsuchida K, Yokoyama M, Inokuchi K (2008) Activin in the brain modulates anxiety-related behavior and adult neurogenesis. PLoSOne 3:e1869.

Aimone JB, Wiles J, Gage FH (2009) Computational influence of adult neurogenesis on memory encoding. Neuron 61:187–202.

Akers KG, Martinez-Canabal A, Restivo L, Yiu AP, De CA, Hsiang HL, Wheeler AL, Guskjolen A, Niibori Y, Shoji H, Ohira K, Richards BA, Miyakawa T, Josselyn SA, Frankland PW (2014) Hippocampal neurogenesis regulates forgetting during adulthood and infancy. Science 344:598–602.

Anon (2016) Handbook of the Psychology of Aging. Elsevier. Available at: https://linkinghub.elsevier.com/retrieve/pii/C20120072213 [Accessed June 19, 2020].

Arredondo SB, Guerrero FG, Herrera-Soto A, Jensen-Flores J, Bustamante DB, Oñate-Ponce A, Henny P, Varas-Godoy M, Inestrosa NC, Varela-Nallar L (2020) Wnt5a promotes differentiation and development of adult-born neurons in the hippocampus by noncanonical Wnt signaling: Wnt5a promotes adult hippocampal neurogenesis. STEM CELLS 38:422–436.

Babayeva S, Zilber Y, Torban E (2011) Planar cell polarity pathway regulates actin rearrangement, cell shape, motility, and nephrin distribution in podocytes. Am J Physiol Renal Physiol 300:F549–560.

Baptista P, Andrade JP (2018) Adult Hippocampal Neurogenesis: Regulation and Possible Functional and Clinical Correlates. Front Neuroanat 12:44.

Berdugo-Vega G, Arias-Gil G, López-Fernández A, Artegiani B, Wasielewska JM, Lee C-C, Lippert MT, Kempermann G, Takagaki K, Calegari F (2020) Increasing neurogenesis refines hippocampal activity rejuvenating navigational learning strategies and contextual memory throughout life. Nat Commun 11:135.

Bergami M, Rimondini R, Santi S, Blum R, Gotz M, Canossa M (2008) Deletion of TrkB in adult progenitors alters newborn neuron integration into hippocampal circuits and increases anxietylike behavior. ProcNatlAcadSciUSA 105:15570–15575.

Bessa JM, Ferreira D, Melo I, Marques F, Cerqueira JJ, Palha JA, Almeida OFX, Sousa N (2009) The mood-improving actions of antidepressants do not depend on neurogenesis but are associated with neuronal remodeling. Mol Psychiatry 14:764–773, 739.

Bunsey M, Eichenbaum H (1996) Conservation of hippocampal memory function in rats and humans. Nature 379:255–257.

Christian KM, Ming G, Song H (2020) Adult neurogenesis and the dentate gyrus: Predicting function from form. Behav Brain Res 379:112346.

David DJ, Samuels BA, Rainer Q, Wang J-W, Marsteller D, Mendez I, Drew M, Craig DA, Guiard BP, Guilloux J-P, Artymyshyn RP, Gardier AM, Gerald C, Antonijevic IA, Leonardo ED, Hen R (2009) Neurogenesis-dependent and-independent effects of fluoxetine in an animal model of anxiety/depression. Neuron 62:479–493.

Davis RL, Zhong Y (2017) The Biology of Forgetting-A Perspective. Neuron 95:490–503.

Dobrossy MD, Drapeau E, Aurousseau C, Le Moal M, Piazza PV, Abrous DN (2003) Differential effects of learning on neurogenesis: learning increases or decreases the number of newly born cells depending on their birth date. MolPsychiatry 8:974–982.

Drapeau E, Abrous DN (2008) Role of neurogenesis in age-related memory disorders. Aging Cell 7:569589.

Drapeau E, Mayo W, Aurousseau C, Le Moal M, Piazza PV, Abrous DN (2003) Spatial memory performances of aged rats in the water maze predict levels of hippocampal neurogenesis. ProcNatlAcadSciUSA 100:14385–14390.

Dupret D, Montaron MF, Drapeau E, Aurousseau C, Le Moal M, Piazza PV, Abrous DN (2005) Methylazoxymethanol acetate does not fully block cell genesis in the young and aged dentate gyrus. EurJNeurosci 22:778–783.

Dupret D, Revest JM, Koehl M, Ichas F, De GF, Costet P, Abrous DN, Piazza PV (2008) Spatial relational memory requires hippocampal adult neurogenesis. PLoSOne 3:e1959.

Eichenbaum H (2000) A cortical-hippocampal system for declarative memory. Nat Rev Neurosci 1:41–50.

Eichenbaum H, Stewart C, Morris RG (1990) Hippocampal representation in place learning. J Neurosci 10:3531–3542.

Eom T-Y, Jope RS (2009) Blocked Inhibitory Serine-Phosphorylation of Glycogen Synthase Kinase-3α/β Impairs In Vivo Neural Precursor Cell Proliferation. Biol Psychiatry 66:494–502.

Epp JR, Silva MR, Kohler S, Josselyn SA, Frankland PW (2016) Neurogenesis-mediated forgetting minimizes proactive interference. Nat Commun 7:10838.

Fiorentini A, Rosi MC, Grossi C, Luccarini I, Casamenti F (2010) Lithium Improves Hippocampal Neurogenesis, Neuropathology and Cognitive Functions in APP Mutant Mice. PLOS ONE 5:e14382.

Gao B, Song H, Bishop K, Elliot G, Garrett L, English MA, Andre P, Robinson J, Sood R, Minami Y, Economides AN, Yang Y (2011) Wnt Signaling Gradients Establish Planar Cell Polarity by Inducing Vangl2 Phosphorylation through Ror2. Dev Cell 20:163–176.

Garthe A, Behr J, Kempermann G (2009) Adult-generated hippocampal neurons allow the flexible use of spatially precise learning strategies. PLoSOne 4:e5464.

Gonzalez-Escamilla G, Muthuraman M, Chirumamilla VC, Vogt J, Groppa S (2018) Brain Networks Reorganization During Maturation and Healthy Aging-Emphases for Resilience. Front Psychiatry 9:601.

Hagiwara A, Yasumura M, Hida Y, Inoue E, Ohtsuka T (2014) The planar cell polarity protein Vangl2 bidirectionally regulates dendritic branching in cultured hippocampal neurons. BioMed Central. Available at: http://molecularbrain.biomedcentral.com/articles/10.1186/s13041-014-0079-5 [Accessed May 27, 2020].

Hill AS, Sahay A, Hen R (2015) Increasing Adult Hippocampal Neurogenesis is Sufficient to Reduce Anxiety and Depression-Like Behaviors. Neuropsychopharmacol Off Publ Am Coll Neuropsychopharmacol 40:2368–2378.

Hirota Y, Sawada M, Kida YS, Huang S, Yamada O, Sakaguchi M, Ogura T, Okano H, Sawamoto K (2012) Roles of Planar Cell Polarity Signaling in Maturation of Neuronal Precursor Cells in the Postnatal Mouse Olfactory Bulb. STEM CELLS 30:1726–1733.

Jang M-H, Bonaguidi MA, Kitabatake Y, Sun J, Song J, Kang E, Jun H, Zhong C, Su Y, Guo JU, Wang MX, Sailor KA, Kim J-Y, Gao Y, Christian KM, Ming G, Song H (2013) Secreted Frizzled-Related Protein 3 Regulates Activity-Dependent Adult Hippocampal Neurogenesis. Cell Stem Cell 12:215–223.

Jessberger S, Clark RE, Broadbent NJ, Clemenson GD Jr, Consiglio A, Lie DC, Squire LR, Gage FH (2009) Dentate gyrus-specific knockdown of adult neurogenesis impairs spatial and object recognition memory in adult rats. LearnMem 16:147–154.

Joly M, Ammersdörfer S, Schmidtke D, Zimmermann E (2014) Touchscreen-based cognitive tasks reveal age-related impairment in a primate aging model, the grey mouse lemur (Microcebus murinus). PloS One 9:e109393.

Kibar Z, Vogan KJ, Groulx N, Justice MJ, Underhill DA, Gros P (2001) Ltap, a mammalian homolog of Drosophila Strabismus/Van Gogh, is altered in the mouse neural tube mutant Loop-tail. Nat Genet 28:251–255.

Lemaire V, Koehl M, Le MM, Abrous DN (2000) Prenatal stress produces learning deficits associated with an inhibition of neurogenesis in the hippocampus. ProcNatlAcadSciUSA 97:11032–11037.

Lie DC, Colamarino SA, Song HJ, Desire L, Mira H, Consiglio A, Lein ES, Jessberger S, Lansford H, Dearie AR, Gage FH (2005) Wnt signalling regulates adult hippocampal neurogenesis. Nature 437:1370–1375.

Liu Y, Du S, Lv L, Lei B, Shi W, Tang Y, Wang L, Zhong Y (2016) Hippocampal Activation of Rac1 Regulates the Forgetting of Object Recognition Memory. Curr Biol CB 26:2351–2357.

Mardones MD, Andaur GA, Varas-Godoy M, Henriquez JF, Salech F, Behrens MI, Couve A, Inestrosa NC, Varela-Nallar L (2016) Frizzled-1 receptor regulates adult hippocampal neurogenesis. Mol Brain 9:29.

Matzel LD, Light KR, Wass C, Colas-Zelin D, Denman-Brice A, Waddel AC, Kolata S (2011) Longitudinal attentional engagement rescues mice from age-related cognitive declines and cognitive inflexibility. Learn Mem Cold Spring Harb N 18:345–356.

McAvoy KM, Sahay A (2017) Targeting Adult Neurogenesis to Optimize Hippocampal Circuits in Aging. Neurotherapeutics 14:630–645.

McAvoy KM, Scobie KN, Berger S, Russo C, Guo N, Decharatanachart P, Vega-Ramirez H, Miake-Lye S, Whalen M, Nelson M, Bergami M, Bartsch D, Hen R, Berninger B, Sahay A (2016) Modulating Neuronal Competition Dynamics in the Dentate Gyrus to Rejuvenate Aging Memory Circuits. Neuron 91:1356–1373.

Montaron M-F, Charrier V, Blin N, Garcia P, Abrous DN (2020) Responsiveness of dentate neurons generated throughout adult life is associated with resilience to cognitive aging. Aging Cell n/a:e13161.

Montaron MF, Drapeau E, Dupret D, Kitchener P, Aurousseau C, Le MM, Piazza PV, Abrous DN (2006) Lifelong corticosterone level determines age-related decline in neurogenesis and memory 22. NeurobiolAging 27:645–654.

Montcouquiol M, Crenshaw EB III, Kelley MW (2006a) Noncanonical Wnt signaling and neural polarity. AnnuRevNeurosci 29:363–386.

Montcouquiol M, Sans N, Huss D, Kach J, Dickman JD, Forge A, Rachel RA, Copeland NG, Jenkins NA, Bogani D, Murdoch J, Warchol ME, Wenthold RJ, Kelley MW (2006b) Asymmetric localization of Vangl2 and Fz3 indicate novel mechanisms for planar cell polarity in mammals. J Neurosci 26:5265–5275.

Mota C, Taipa R, das Neves SP, Monteiro-Martins S, Monteiro S, Palha JA, Sousa N, Sousa JC, Cerqueira JJ (2019) Structural and molecular correlates of cognitive aging in the rat. Sci Rep 9:2005.

Munji RN, Choe Y, Li G, Siegenthaler JA, Pleasure SJ (2011) Wnt signaling regulates neuronal differentiation of cortical intermediate progenitors. J Neurosci Off J Soc Neurosci 31:1676–1687.

Murdoch JN (2001) Severe neural tube defects in the loop-tail mouse result from mutation of Lpp1, a novel gene involved in floor plate specification. Hum Mol Genet 10:2593–2601.

Nyberg L, Pudas S (2019) Successful Memory Aging. Annu Rev Psychol 70:219–243.

Okerlund ND, Stanley RE, Cheyette BNR (2016) The Planar Cell Polarity Transmembrane Protein Vangl2 Promotes Dendrite, Spine and Glutamatergic Synapse Formation in the Mammalian Forebrain. Mol Neuropsychiatry 2:107–114.

Palomer E, Buechler J, Salinas PC (2019) Wnt Signaling Deregulation in the Aging and Alzheimer’s Brain. Front Cell Neurosci 13:227.

Raber J, Rola R, LeFevour A, Morhardt D, Curley J, Mizumatsu S, VandenBerg SR, Fike JR (2004) Radiation-induced cognitive impairments are associated with changes in indicators of hippocampal neurogenesis. RadiatRes 162:39–47.

Revest JM, Dupret D, Koehl M, Funk-Reiter C, Grosjean N, Piazza PV, Abrous DN (2009) Adult hippocampal neurogenesis is involved in anxiety-related behaviors. MolPsychiatry 14:959–967.

Robert BJA, Moreau MM, Dos Santos Carvalho S, Barthet G, Racca C, Bhouri M, Quiedeville A, Garret M, Atchama B, Al Abed AS, Guette C, Henderson DJ, Desmedt A, Mulle C, Marighetto A, Montcouquiol M, Sans N (2020) Vangl2 in the Dentate Network Modulates Pattern Separation and Pattern Completion. Cell Rep 31:107743.

Ross AJ et al. (2005) Disruption of Bardet-Biedl syndrome ciliary proteins perturbs planar cell polarity in vertebrates. NatGenet 37:1135–1140.

Santarelli L, Saxe M, Gross C, Surget A, Battaglia F, Dulawa S, Weisstaub N, Lee J, Duman R, Arancio O, Belzung C, Hen R (2003) Requirement of hippocampal neurogenesis for the behavioral effects of antidepressants. Science 301:805–809.

Saxe MD, Battaglia F, Wang JW, Malleret G, David DJ, Monckton JE, Garcia AD, Sofroniew MV, Kandel ER, Santarelli L, Hen R, Drew MR (2006) Ablation of hippocampal neurogenesis impairs contextual fear conditioning and synaptic plasticity in the dentate gyrus. ProcNatlAcadSciUSA 103:17501–17506.

Schafer ST, Han J, Pena M, von Bohlen und Halbach O, Peters J, Gage FH (2015) The Wnt Adaptor Protein ATP6AP2 Regulates Multiple Stages of Adult Hippocampal Neurogenesis. J Neurosci 35:4983–4998.

Seib DRM, Corsini NS, Ellwanger K, Plaas C, Mateos A, Pitzer C, Niehrs C, Celikel T, Martin-Villalba A (2013) Loss of Dickkopf-1 Restores Neurogenesis in Old Age and Counteracts Cognitive Decline. Cell Stem Cell 12:204–214.

Shors TJ, Townsend DA, Zhao M, Kozorovitskiy Y, Gould E (2002) Neurogenesis may relate to some but not all types of hippocampal-dependent learning. Hippocampus 12:578–584.

Sun J, Bonaguidi MA, Jun H, Guo JU, Sun GJ, Will B, Yang Z, Jang M-H, Song H, Ming G, Christian KM (2015) A septo-temporal molecular gradient of sfrp3 in the dentate gyrus differentially regulates quiescent adult hippocampal neural stem cell activation. Mol Brain 8:1–10.

Surget A, Tanti A, Leonardo ED, Laugeray A, Rainer Q, Touma C, Palme R, Griebel G, Ibarguen-Vargas Y, Hen R, Belzung C (2011) Antidepressants recruit new neurons to improve stress response regulation. Mol Psychiatry 16:1177–1188.

Thakar S, Wang L, Yu T, Ye M, Onishi K, Scott J, Qi J, Fernandes C, Han X, Yates JR, Berg DK, Zou Y (2017) Evidence for opposing roles of Celsr3 and Vangl2 in glutamatergic synapse formation. Proc Natl Acad Sci 114:E610–E618.

Tissir F, Bar I, Jossin Y, De BO, Goffinet AM (2005) Protocadherin Celsr3 is crucial in axonal tract development. NatNeurosci 8:451–457.

Tissir F, Goffinet AM (2006) Expression of planar cell polarity genes during development of the mouse CNS. EurJNeurosci 23:597–607.

Tissir F, Goffinet AM (2013) Shaping the nervous system: role of the core planar cell polarity genes. Nat Rev Neurosci 14:525–535.

Tran LM, Josselyn SA, Richards BA, Frankland PW (2019) Forgetting at biologically realistic levels of neurogenesis in a large-scale hippocampal model. BehavBrain Res 376:112180.

van Boxtel MP, Buntinx F, Houx PJ, Metsemakers JF, Knottnerus A, Jolles J (1998) The relation between morbidity and cognitive performance in a normal aging population. J Gerontol A Biol Sci Med Sci 53:M147–154.

Varela-Nallar L, Inestrosa NC (2013) Wnt signaling in the regulation of adult hippocampal neurogenesis. Front Cell Neurosci 7 Available at: http://journal.frontiersin.org/article/10.3389/fncel.2013.00100/abstract [Accessed April 1, 2020].

Wada H, Iwasaki M, Sato T, Masai I, Nishiwaki Y, Tanaka H, Sato A, Nojima Y, Okamoto H (2005) Dual roles of zygotic and maternal Scribble1 in neural migration and convergent extension movements in zebrafish embryos. Development 132:2273–2285.

Wang Y, Zhang J, Mori S, Nathans J (2006) Axonal growth and guidance defects in Frizzled3 knock-out mice: a comparison of diffusion tensor magnetic resonance imaging, neurofilament staining, and genetically directed cell labeling. J Neurosci 26:355–364.

Wexler EM, Paucer A, Kornblum HI, Palmer TD, Geschwind DH (2009) Endogenous Wnt Signaling Maintains Neural Progenitor Cell Potency. STEM CELLS 27:1130–1141.

Yang W, Garrett L, Feng D, Elliott G, Liu X, Wang N, Wong YM, Choi NT, Yang Y, Gao B (2017) Wnt-induced Vangl2 phosphorylation is dose-dependently required for planar cell polarity in mammalian development. Cell Res 27:1466–1484.

Yang W, Zhou X, Ma T (2019) Memory Decline and Behavioral Inflexibility in Aged Mice Are Correlated With Dysregulation of Protein Synthesis Capacity. Front Aging Neurosci 11:246.

Yin H, Copley CO, Goodrich LV, Deans MR (2012) Comparison of Phenotypes between Different vangl2 Mutants Demonstrates Dominant Effects of the Looptail Mutation during Hair Cell Development. PLOS ONE 7:e31988.

